# The Role of a Crystallographically Unresolved Cytoplasmic Loop in Stabilizing the Bacterial Membrane Insertase YidC2

**DOI:** 10.1101/707778

**Authors:** Thomas Harkey, Vivek Govind Kumar, Jeevapani Hettige, Seyed Hamid Tabari, Kalyan Immadisetty, Mahmoud Moradi

## Abstract

YidC, a bacterial member of the YidC/Alb3/Oxa1 insertase family, mediates membrane protein assembly and insertion. Cytoplasmic loops are known to have functional significance in membrane proteins such as YidC. Employing microsecond-level molecular dynamics (MD) simulations, we show that the crystallographically unresolved C2 loop plays a crucial role in the structural dynamics of *Bacillus halodurans* YidC2. We have modeled the C2 loop and used allatom MD simulations to investigate the structural dynamics of YidC2 in its *apo* form, both with and without the C2 loop. The C2 loop was found to stabilize the entire protein and particularly the C1 region. C2 was also found to stabilize the alpha-helical character of the C-terminal region. Interestingly, the highly polar or charged lipid head groups of the simulated membranes were found to interact with and stabilize the C2 loop. These findings demonstrate that the crystallographically unresolved loops of membrane proteins could be important for the stabilization of the protein despite the apparent lack of structure, which could be due to the absence of the relevant lipids to stabilize them in crystallographic conditions.

## INTRODUCTION

One third of all genes encode membrane proteins, which must be folded and inserted into the plasma membrane co-translationally^1,2^. The YidC/Oxa1/Alb3 family of membrane proteins mediates the proper folding and insertion of incoming peptides and proteins in the membrane^3,4,5,6^. Mammalian Oxa1 (found in mitochondria), plant Alb3 (found in chloroplasts), and YidC (found in bacteria) are homologous insertases^5^. Insertases exist in all domains of life and are essential for the viability of cells^7,8,9^. They are able to function either in a Sec-dependent or a Sec-independent manner^10,11,12,13,14,15,16,17^. In this study, we investigate the structural dynamics of YidC, which is the most well characterized member of the family.

Several studies have investigated the role of the YidC protein in different organisms. YidC plays an important role in folding of the LacY lactose permease membrane protein and is essential for the insertion of the *c* subunit of the F_0_F_1_-ATPase (F0c) into the plasma membrane of *Escherichia coli* (a gram-negative bacterium) in a Sec-independent manner^11,14,18^. The genomes of most gram-positive bacteria encode two YidC proteins, YidC1 and YidC2^19^. Both paralogs have functional overlap but YidC2 may have a function not shared by YidC1^20^. Multiple crystal structures of YidC are available. Kumazaki et al. have crystallized YidC2 from the gram-positive bacterium *Bacillus halodurans* and proposed a binding and insertion mechanism for single-spanning membrane proteins^5^. Here, we investigate the structural dynamics of functionally important regions of YidC2 from *Bacillus halodurans* (PDB entry: 3WO7)^5^.

YidC2 consists of five transmembrane (TM) helices (TM1-5) connected by two cytoplasmic regions (C1 and C2) and two extracellular regions (E2 and E3)^5^. The C1 region consists of two helices (CH1 and CH2) connected by a short loop^21^. The insertase function appears to be localized to the TM region^22^, with TM3 being most crucial for function^23^. Yuan et al., have investigated mutations in the TM3 segment of YidC (C423R and P431L)^23^. C423R has been found to produce a weak membrane protein insertion phenotype while P431L results in a stronger insertion phenotype^23^. The Pf3 coat protein and the *c* subunit of the F_0_F_1_-ATPase are the affected substrates in both cases^23^. Several hydrophilic residues, including a positively charged arginine at position 72 (R72), form a groove that is accessible to both the cytoplasm and the lipid bilayer^5^. Figure 1 shows the position of these different domains.

**Figure 1.**
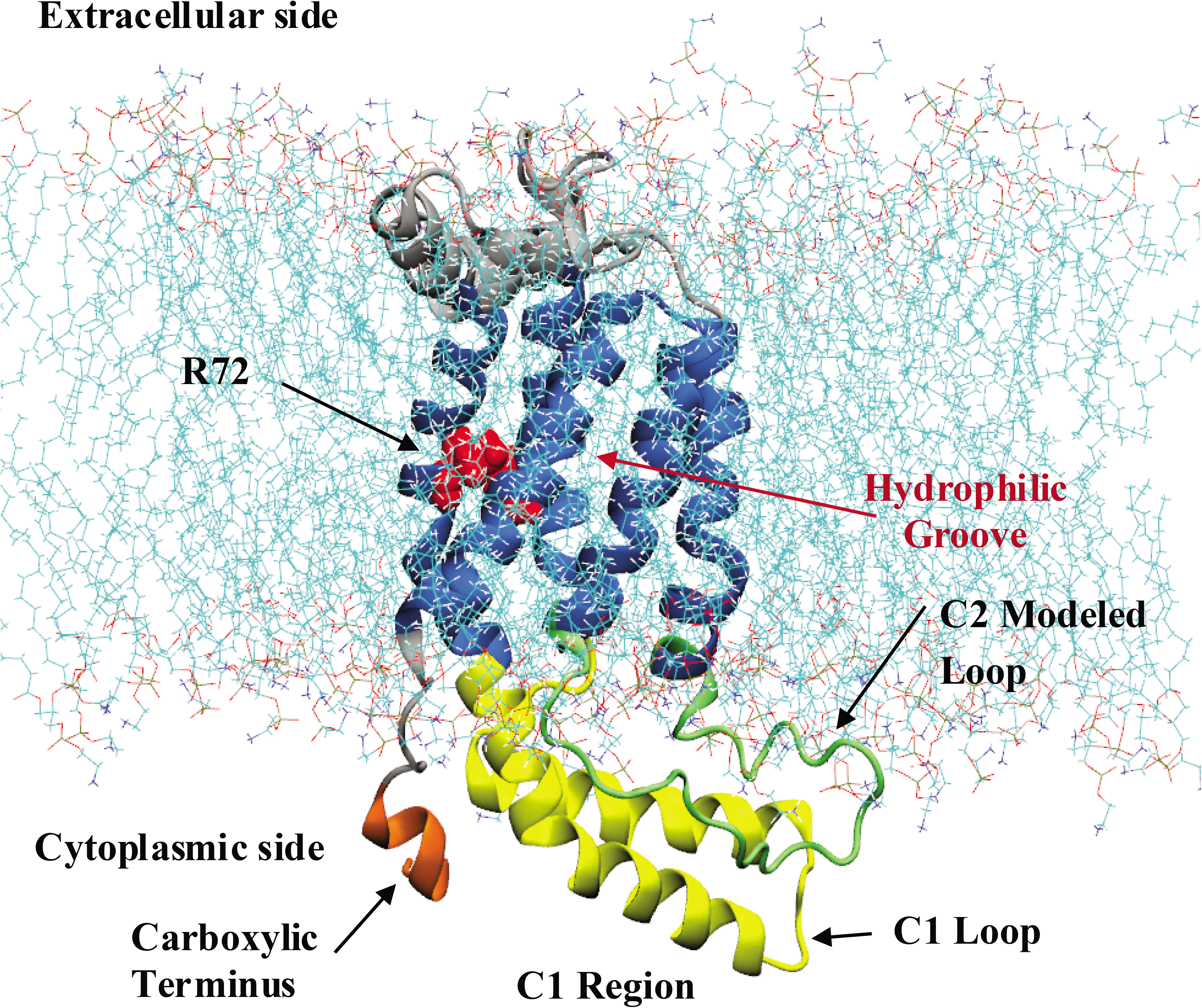
The bacterial membrane protein insertase YidC2. The cartoon representation of membrane embedded YidC2 model based on its crystal structure (<u>**PDB ID: 3WO7**</u>)(Kumazaki et al., 2014a) is shown. The five TM helices (blue) of YidC2, the hydrophilic groove, the side chain of residue R72 (red), the C1 region including CH1 and CH2 helices connected with the C1 loop (yellow), the modeled C2 loop (green), and the carboxyl terminal helix (orange) are shown. The modeled cytoplasmic loop C2 (residues 195-216) was not resolved in the crystal structure.

Kumazaki *et al.*^5^ have hypothesized that the incoming substrate first interacts with the C1 loop, and is subsequently netted into the hydrophilic groove of YidC, with negatively charged residues on the incoming substrate interacting with the positively charged R72 in the groove region^5,24^. The importance of a conserved arginine residue in the hydrophilic groove has also been reported for *E. coli* YidC^25^. High B-factor values have been reported for the C1 region, indicating that this region fluctuates greatly^5^. MD simulations also agree with the assessment that the C1 region is flexible^5^. In another study, Kumazaki *et al.* report that the C1 region is flexible in *E. coli* YidC (based on B-factor values), suggesting that the C1 region flexibility is a universal characteristic of YidC^21^. The C-terminal domain is also thought to have functional relevance and may be involved in the folding of the periplasmic regions of inserted substrates^26^. The C2 loop is not resolved in any of the YidC2 crystal structures^5^. The C2 loop of *E. coli* YidC was recently resolved in an *E. coli* YidC crystal structure^27^ and suggested to play a role in the activation mechanism of YidC by covering the hydrophilic groove in its inactive state (as in the resolved structure) and by exposing the hydrophilic groove upon activation triggered by ribosome^27^. However, the C2 loop of *E. coli* YidC is considerably shorter than that of YidC2 in gram-positive bacteria, and the two may have different functional roles due to the significant difference in their length.

The importance of cyto- and periplasmic loops for the functioning of membrane proteins has been studied extensively in various proteins such as NS4B^28^, Wzy^29^, and LolCDE^30^. Due to the importance of cytoplasmic loops in membrane proteins, we have modeled the crystallographically unresolved C2 loop of YidC2 in order to investigate the impact of this loop on overall protein stability as well as the stability of functionally important regions such as the C1 loop and the C-terminal domain. All-atom MD provides a reliable method for the investigation of membrane protein structural dynamics particularly in a comparative manner^31^. Nanosecond-level MD simulations, however, as are often used for such studies, have been shown to be questionable for reliably describing the functionally relevant conformational dynamics of membrane proteins^32^. Therefore, either microsecond-level^31^ and/or enhanced sampling^33,34,35^ simulations must be employed to investigate the conformational behavior of membrane proteins. Here we have employed microsecond-level all-atom MD simulations of membrane-embedded YidC2 that show a key role for the C2 cytoplasmic loop in the structural dynamics of YidC2.

## RESULTS AND DISCUSSION

We have performed several sets of unbiased all-atom MD simulations of YidC2 to examine the importance of the crystallographically unresolved C2 loop of YidC2. To simplify the comparison, we first conducted the simulations of YidC2 with and without the C2 loop in a pure 1-palmitoyl-2-oleoyl-sn-glycero-3-phosphoethanolamine (POPE) environment. The initial models are both based on the crystal structure of YidC2 (PDB entry: 3WO7)^5^ in the presence of explicit membrane and water. The missing C2 loop was modeled carefully in one of the simulations and was not present in the other (see Methods for simulation details). Since the two systems are virtually identical besides the presence/absence of the C2 loop, we can make meaningful comparisons between the two sets of simulations. Each simulation was performed for 2 microseconds, which is long enough to reveal meaningful differences between the two sets. Any difference that is observed in the flexibility of the protein in each system can be attributed to the presence or absence of the C2 loop, given that the two trajectories are converged. We have also performed control simulations in a mixed lipidic environment and with an alternative C2 loop model to ensure our major findings are reproducible. We first focus on the results from the two 2-μs long simulations of YidC2 in POPE membrane.

We have identified several regions of interest in YidC2 to characterize the conformational dynamics of both systems. Figure 1 identifies some of these regions in the system with the modeled C2 loop. These regions include the C1 region, the C2 modeled loop, the TM helices, and the carboxyl terminal region. The C1 region (including the C1 loop and CH1/CH2 helices), carboxyl terminal region, and C2 loop all reside in the cytoplasmic area.

### The presence of the C2 loop stabilizes the global protein conformation

Comparing the *C_α_* root mean square deviation (RMSD) of the two systems indicates that the presence of the C2 loop stabilizes the protein (Figure 2A). The RMSD of YidC2 was calculated and plotted as a function of simulation time. The system without the loop is clearly less stable (RMSD=3.0±0.41 Å) than the system with the loop (RMSD=2.4±0.40 Å) (Figure 2A). Intriguingly, root mean square fluctuation (RMSF) analysis indicates that the stabilization is not localized but a large part of the protein shows a more rigid conformation in the presence of the C2 loop, evidenced by a lower value of RMSF for most residues (Figure 2B). The difference in the RMSF values, however, is not the same for all domains, indicating that some domains are more directly influenced by the presence of the C2 loop. Overall, the protein shows a relatively dynamic behavior in both cases (Figure 2A), however, our results indicate that the flexibility of the protein is considerably exaggerated when the C2 loop is not considered, as revealed more clearly using the principal component analysis (PCA).

**Figure 2.**
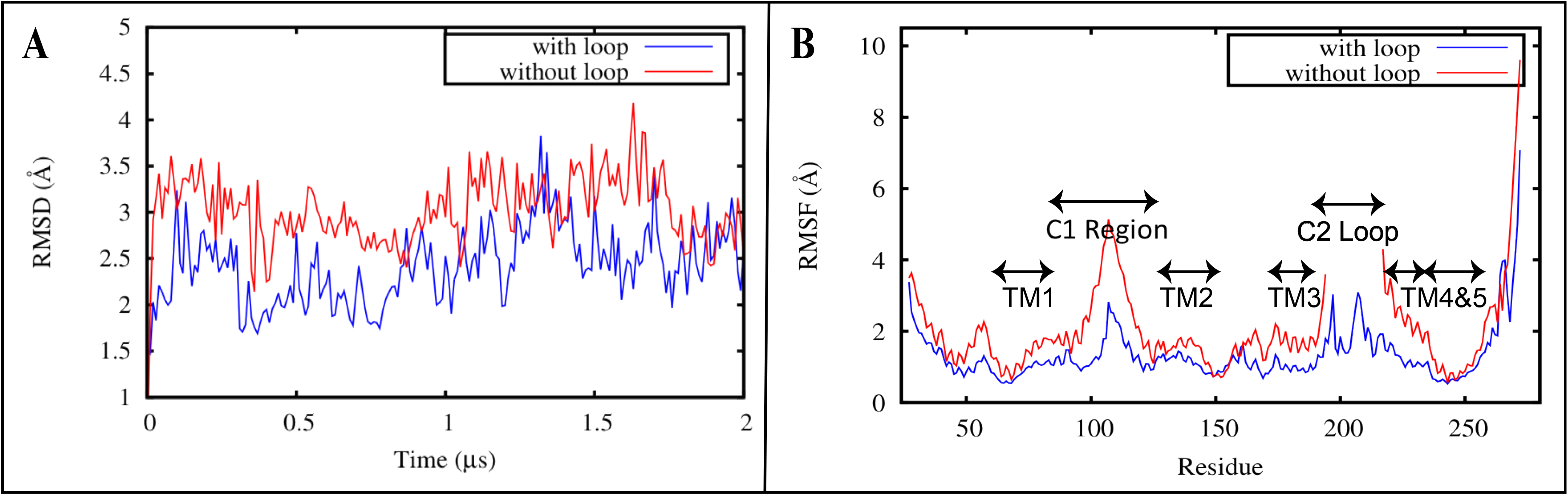
Protein stability assessment for YidC2 with and without the C2 loop. **A** Root mean square deviation (RMSD) time series and **B** root mean square fluctuation (RMSF) estimations for YidC2 with (blue) and without (red) the C2 loop obtained from 2-μs simulations. RMSD analysis indicates that the absence of the C2 loop destabilizes the protein and RMSF analysis indicates that the presence of the modeled C2 loop (residues 195 to 216) significantly stabilizes the entire protein, particularly in the C1 loop region (residues 84-133).

We performed PCA to verify our claim that the C2 loop stabilizes the protein (Figure 3A). When both of the trajectories are projected onto the space of their first two principal components (PC1 and PC2), it clearly demonstrates that the system with the loop clusters tightly around a specific region as compared to the system without the loop that samples a number of scattered regions in the configuration space. This clearly indicates that the system with the loop is more stable than the system without the loop. For a discussion on the convergence of the PCA calculations see Supplemental Figure S1, showing the results of PCA based on the first half of the trajectories only, which agrees with the results shown in Figure 3 based on the entire trajectories.

**Figure 3.**
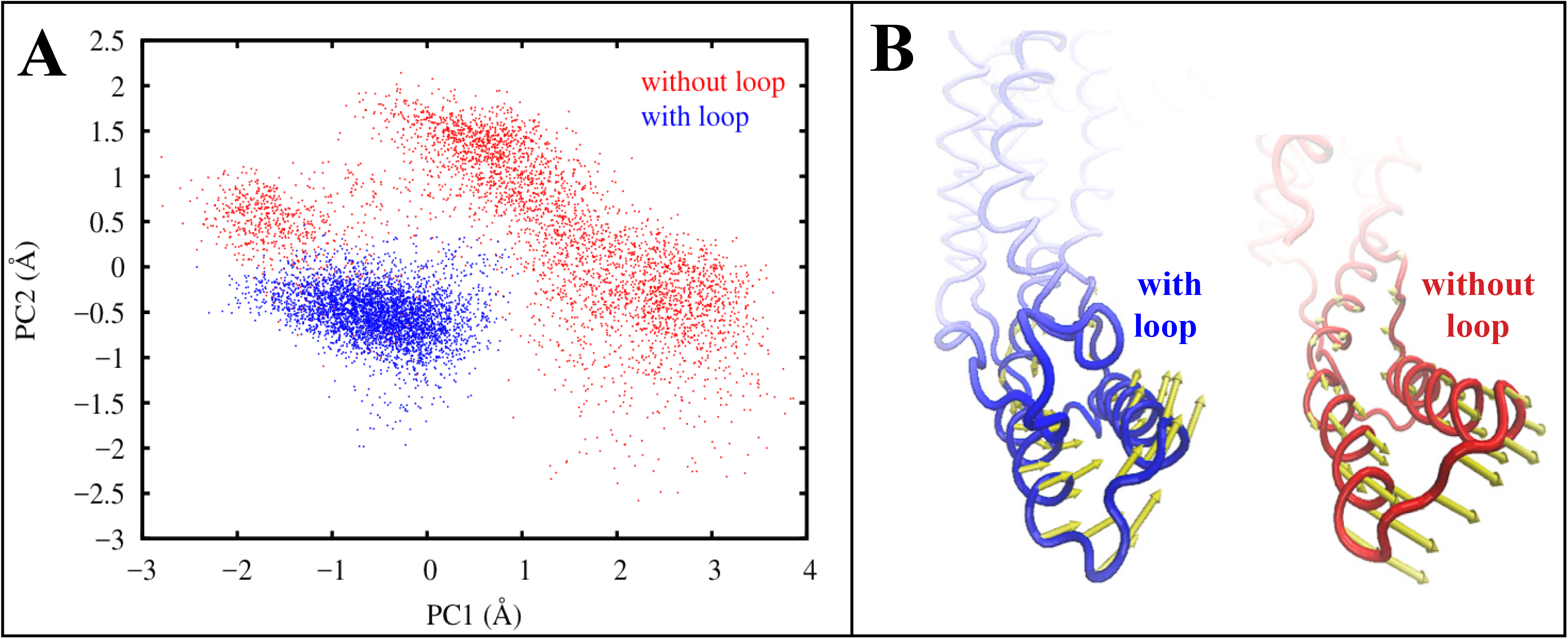
Principal component analysis demonstrates that the system with the C2 loop is more stable than the system without the loop. **A** Projection of the simulation trajectory of YidC2 with (blue) and without (red) the C2 loop onto their first two principal components (PC1, PC2). The YidC2 model with the loop is quite stable only locally fluctuating around its average structure; however, the YidC2 model without the loop jumps between multiple conformations indicating a significant conformational flexibility. **B** First principal component (PC1) eigenvector shown for the C1 region, that shows substantially different behavior in the presence (blue) and absence (red) of the C2 loop. The width of the arrows represents the magnitude of fluctuation.

### The C2 loop increases the stability of the C1 loop

We observe that in the system with the C2 loop, the C1 region is significantly more stable than it is in the system without the C2 loop (Figure 2B). Since the only difference between the two systems is the presence of the C2 loop, we conclude that the presence of the C2 loop stabilizes the C1 region of YidC2. RMSD analysis of the C1 region confirms that the difference in C1 fluctuations is due to the presence of the C2 loop. We calculated the overall and internal RMSD of the C1 loop (Supplemental Figure S2). The former reflects the internal conformational changes plus the translational and rotational motions, while the latter only reflects the internal conformational changes. The overall RMSD for the C1 region without the C2 loop (3.5±0.90 Å) is greater than that with the C2 loop (2.9±0.61 Å) (Supplemental Figure S2A), confirming that the presence of the C2 loop stabilizes the C1 region of YidC. However, there is no significant difference in the behavior of the internal RMSD of the two systems (1.6±0.33Å vs. 1.7±0.30 Å) (Supplemental Figure S2B). A comparison of the internal and overall RMSDs of the C1 region indicates that while the internal conformation of the C1 region stays close to that captured in the crystal structure of YidC2, its orientation does not stay the same in the two cases. PCA analysis illustrates this observation more clearly. The most significant protein collective motion (represented by PC1) in each system is associated with a distinct motion in the C1 region (Figure 3B), which clearly illustrates a difference in behavior of the C1 region between the two systems. The C1 region moves upward towards the C2 loop and becomes stable when the C2 loop is present. However, in the absence of the C2 loop, the C1 region can move downward away from the membrane and fluctuate strongly.

### Salt bridge interactions play a key role in the stabilization of the C1 region by the C2 loop

The difference in the flexibility of YidC2 with and without the C2 loop is particularly pronounced in the C1 region as discussed above. This is due, to a great extent, to the electrostatic interactions between the C1 region and C2 loop. The negatively charged side chain of D205 in the C2 loop can specifically interact with K104 and K109 of the C1 loop (Figure 4A), although the interactions are stronger and more frequent between D205 and K109, as reflected in the salt bridge distance time series (Figure 4B). This is expected considering that D205 is closer to K109 than K104. These salt bridges cannot form in the system without the C2 loop. Therefore, we propose that these salt bridges play a key role in stabilizing the C1 region in the presence of the C2 loop and that in the system without the C2 loop, the C1 region is anchored less strongly to the rest of the protein and is able to fluctuate in the cytoplasm more freely.

**Figure 4:**
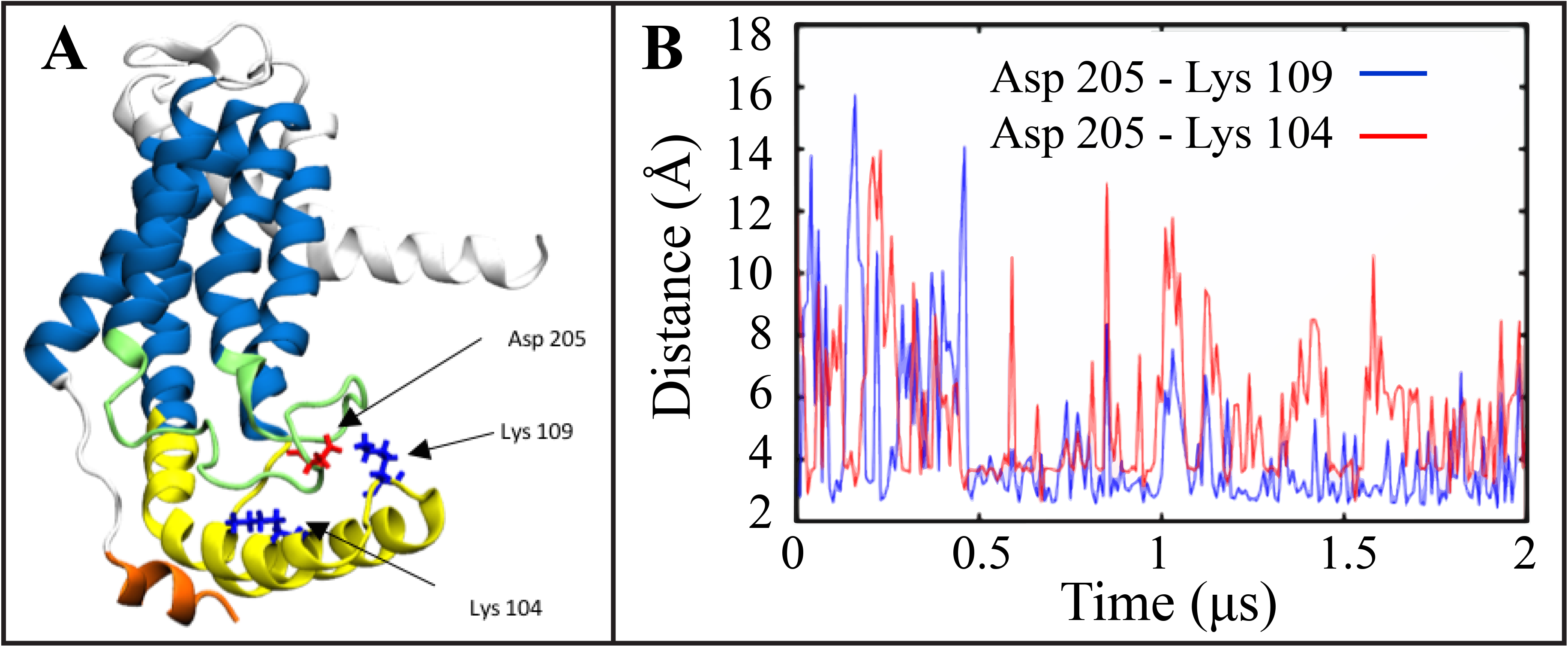
A salt bridge interaction network stabilizes the C1 loop in the presence of the C2 loop. **A** D205 (red) of C2 loop (green) can potentially form a salt bridge with K109 and/or K104 (blue) of the C1 region (yellow). **B** Time series of the D205-K109/104 donor-acceptor salt bridge distances.

We also identified an intradomain salt bridge in the CH1 region of the C1 loop between R93 and E97. The E97-R93 salt bridge distance was significantly lower in the system with the C2 loop (4.3±0.64 Å) (Supplemental Figure S3A,B) than the system without the loop (5.6±1.7 Å) (Supplemental Figure S3C,D). We hypothesize that this salt bridge is destabilized in the system without the loop due to the fluctuating behavior of the C1 region in the absence of the C2 loop. These findings clearly indicate a relationship between the functionally important C1 region and the neighboring C2 loop in the cytoplasmic side of YidC2. Our findings are in correlation with those of Geng *et al.^26^*, where they studied the C2 mediated YidC-ribosome binding in *E. coli* and determined that the C2 region of YidC was involved in ribosome binding, but not the C1 region. They proposed that the C1 region may be involved in downstream activity but not ribosome binding. They also found that the positively charged C-terminus of YidC does play a significant role in the ribosome binding.

### Carboxyl-terminal domain stability and conformation

The C-terminal domain of YidC has been proposed to play a significant role in ribosome binding^26^. Due to its potential functional importance, we probed the conformational dynamics of the C-terminal region of YidC2. The available crystal structure of YidC2^5^ has a modified C-terminal domain with several missing and mutated residues (see Methods). However, we observed a significant difference in its secondary structure in the absence and presence of the C2 loop, an observation that could indicate a potential allosteric interaction between the C-terminal domain and the C2 loop. Over the course of the simulation without the C2 loop, the C-terminal region appears to unravel into a random coil (Figure 5) as evidenced by the secondary structure analysis, where it completely loses its *α*-helical character within the first 0.3 μs of simulation (Figure 5C). On the other hand, the C-terminal region exists as an *α*-helix for the majority of the simulation in the system with the loop (Figure 5A,B). This indicates that the C2 loop stabilizes the C-terminal *α*-helix.

**Figure 5.**
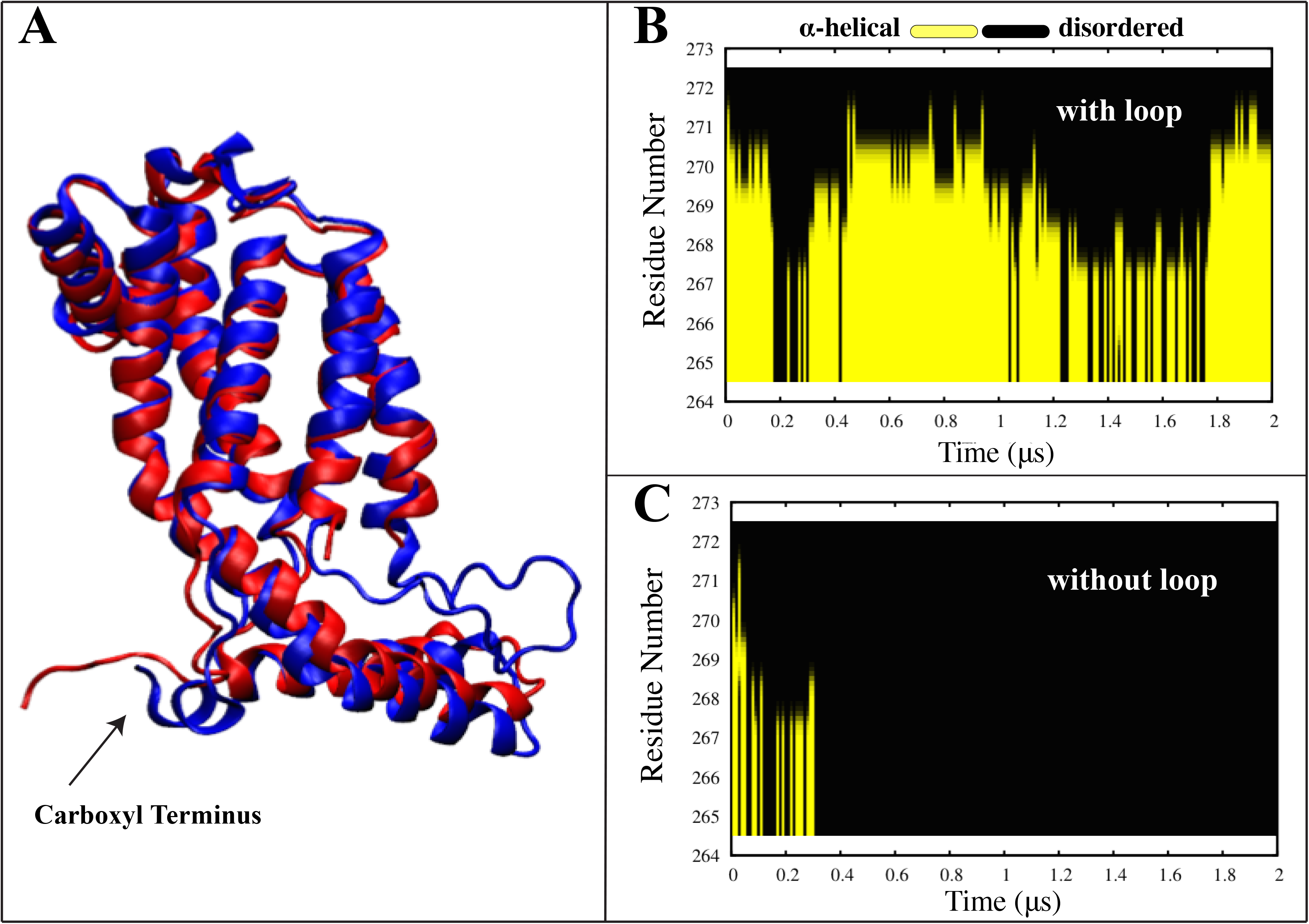
Carboxyl terminal secondary structure. **A** Visual representation of the carboxyl terminal domain of the systems with the loop (blue) and without the loop (red) upon equilibration. α-helical character of the residues 265 to 272 of the modified carboxyl terminal domain with (**B**) and without (**C**) the C2 loop is shown as a function of the simulation time. Yellow color indicates an alpha helical secondary structure. The carboxyl terminal helix unravels into a coil in the system without the loop, whereas it retains the secondary structure for the entire simulation in the system with the loop.

We also analyzed the overall and internal RMSD of the C-terminal domain to characterize its flexibility. The presence of the C2 loop corresponds to a more stable C-terminal domain (overall RMSD = 4.7±1.6 Å) as compared to the model without the C2 loop (overall RMSD = 7.4±1.9 Å) (Supplemental Figure S4A). The internal RMSD for the systems with and without the loop are 2.0±0.81 Å and 3.6±0.62 Å, respectively (Supplemental Figure S4B), demonstrating that there is a difference in internal conformation of the carboxyl terminus regions between the two systems, as also evidenced by our secondary structure analysis. These observations confirm the role of the C2 loop in promoting the helical structure of the C-terminal domain.

### C2 loop allosterically impacts the behavior of other functionally important regions of YidC2

Thus far, we have shown that the C2 loop directly interacts with the C1 region and changes its behavior, while the C-terminal region is also influenced by the presence or absence of the C2 loop. The C-terminal region, however, does not interact directly with the C2 loop. Instead, the C2 loop is affecting the behavior of the C-terminal region indirectly through the C1 region as we will discuss in more detail below.

To systematically investigate the allosteric interactions of C2 loop with different protein domains, we employed dynamic network analysis^36^, which characterizes the linear correlation between different residue pairs. Figure 6 shows the correlation coefficient of each residue pair calculated from the trajectory with the C2 loop, subtracted from the same quantity calculated from the trajectory without the loop, and reported as its absolute value. The reported quantity for each residue pair quantifies the magnitude of the difference in the correlation behavior of the two residues caused by the C2 loop. We observe that the introduction or removal of the C2 loop leads to significant changes in the correlations of different domains of YidC2. Specifically, differences in inter-domain correlations between TM1/C1 region and TM3/TM4 region as well as intradomain correlations of the TM1 helix are quite significant (Fig. 6). However, the specific residues that exhibit the most significant C2-dependent behavior in their correlations with other residues are located in the carboxyl tail that interact strongly with various residues in the C1 region and TM4 helix (Supplemental Figure S5).

**Figure 6.**
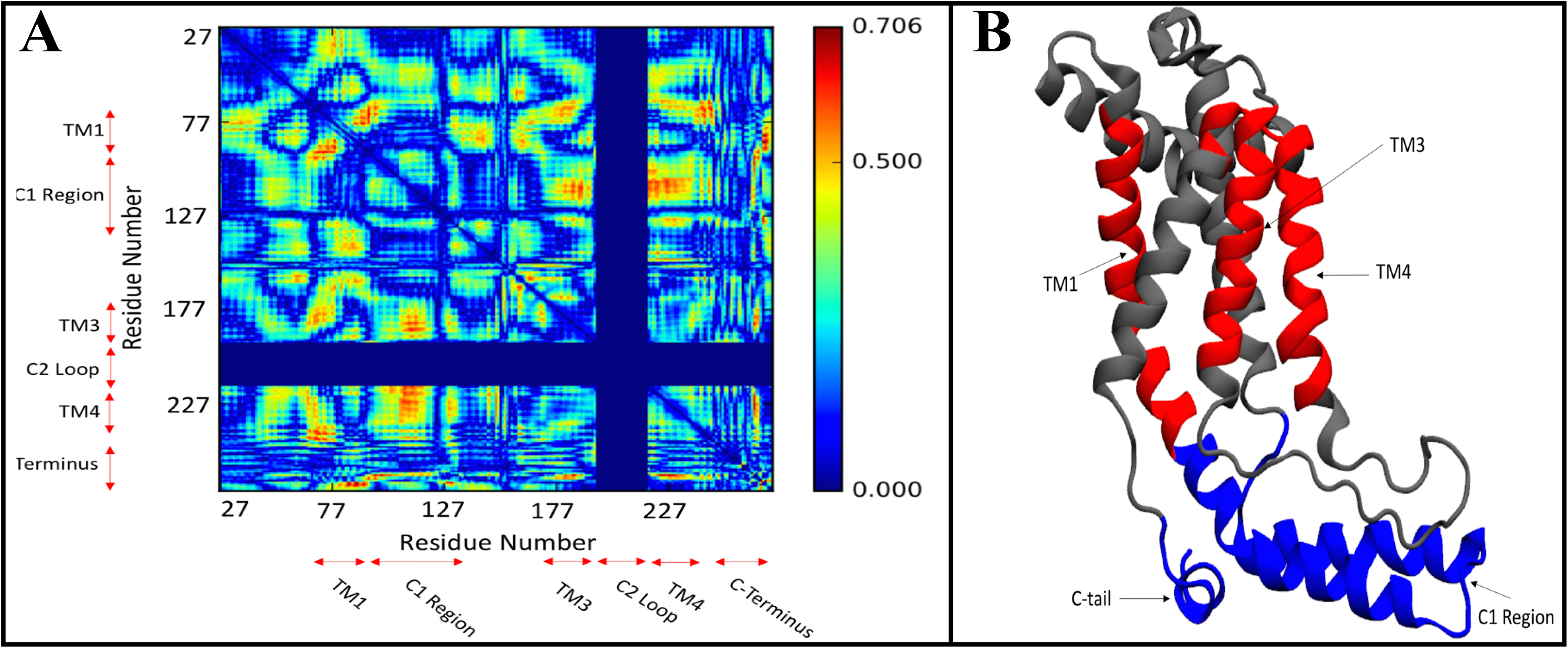
Dynamic network analysis of the systems with and without the C2 loop. **A** The heat map of the absolute value of the residue pair correlation difference between the systems with and without the C2 loop. The x and y axes represent the residue numbers for each residue pair. The blue color indicates no significant change in residue pair correlations due to the presence/absence of the C2 loop. The red color indicates a significant change in the residue pair correlations due to the presence/absence of the C2 loop. **B** The regions with the most significant change in their correlation behavior including TM1, TM3, and TM4 (red) in the transmembrane domain and the C-terminal tail and the C1 region (blue) in the cytoplasmic side are highlighted.

The influence of C2 loop on the C-terminal tail is clear from the analyses above and since there is no direct interaction between the two domains, it is likely that the C1 region mediates the change in the conformation and interactions of the C-terminal domain. For instance, a salt bridge can form between E266 (C-terminal region) and K81 (C1 region) in the system without the C2 loop, which is completely absent in the system with the C2 loop (Figure 7). In the absence of the C2 loop, the C1 region can more freely move and interact with the C-terminal tail. The attraction between the E266 and K81 residues can pull the C-terminal close to the C1-TM1 region. The interaction between the C1 region and the C-terminal domain is likely to be the cause of the disruption of its helical structure, accounting for the difference in C-terminal flexibility between the two systems. E266 in the C-terminal tail is able to move closer to K81 due to the fact that the C1 region, TM4, and TM5 helices of the protein are not interacting/connected to the missing C2 loop. Overall, these findings indicate that the presence of the C2 loop influences the behavior of the functionally important regions of YidC2.

**Figure 7:**
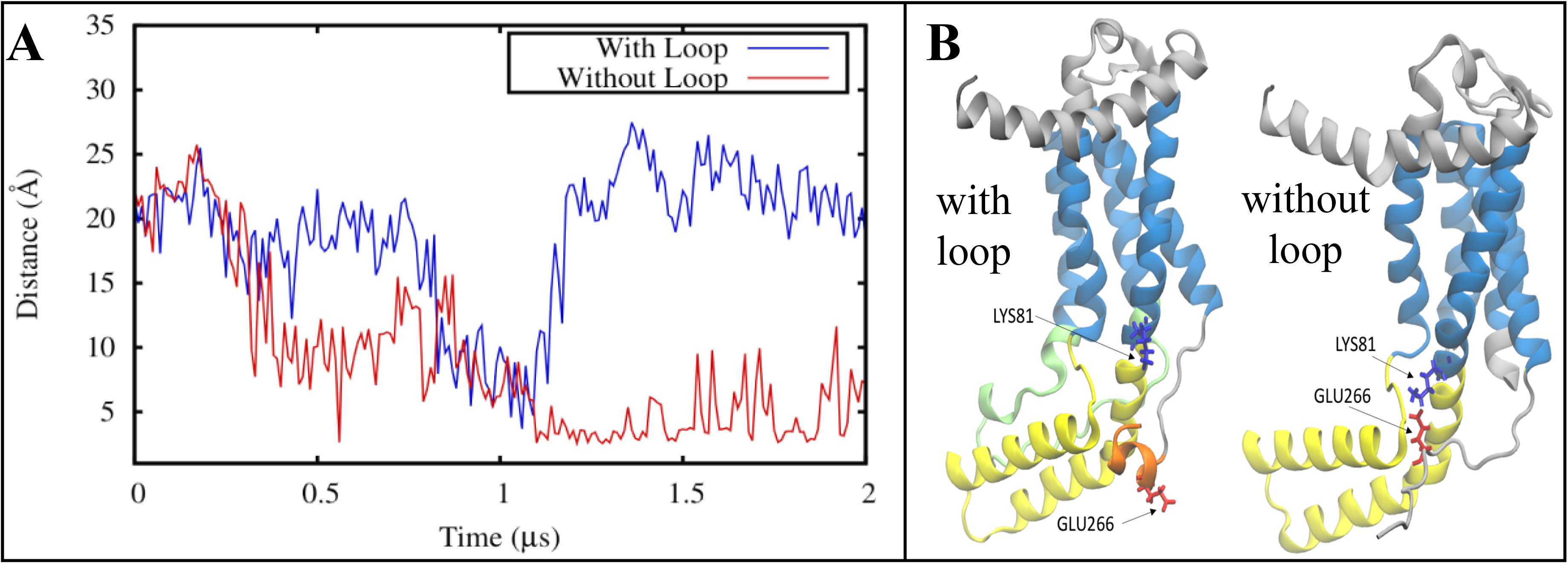
An interdomain salt bridge forms between the C-terminal tail and the C1 region in the absence of the C2 loop. **A** Donor-acceptor salt bridge distance between C-terminal residue E266 and C1 residue K81 as a function of simulation time for the systems without (red) and with (blue) the C2 loop. **B** A visual representation of the K81-E266 interaction in the system with (left) and without the C2 loop (right).

### Interactions with the membrane contributes to the stability of the C2 loop

The absence of the C2 loop not only results in some local conformational changes in YidC2, as discussed above, but also induces global changes. Interestingly, the protein is placed in the membrane somewhat differently in the absence and presence of the C2 loop. Figure 8A illustrates how the tilt angle of the TM region of protein with respect to the membrane normal is distributed differently when the C2 loop is present or absent. The difference in tilt angle is most likely due to the C2 loop membrane interactions. The presence of the C2 loop changes the interaction pattern of the protein with the membrane as shown in Figure 8B. Due to its close proximity to the membrane, the C2 loop interacts with membrane, particularly between the side chains of the C2 loop and the lipid head groups. Although the presence of the C2 loop introduces new interactions between the protein and the membrane, it reduces the total number of lipids interacting with the protein (Figure 8B) by partially blocking other regions that otherwise interact with the membrane (Supplementary Figure S6). Despite the reduction in the number of interacting lipids, however, the net effect is having a more stable protein when the C2 loop is present as previously discussed. This is due to the strong interactions between the C2 loop and the membrane. Specifically, residue D207 of the C2 loop forms one or more hydrogen bonds with the lipid head groups throughout the simulation (Figure 8C). Our interaction energy analysis indicates that the C2-membrane interactions, which are predominantly electrostatic, are steady throughout the simulation (Figure 8D). The interactions between the POPE head groups and the C2 loop provide a possible mechanism for the stabilization of the C2 loop.

**Figure 8.**
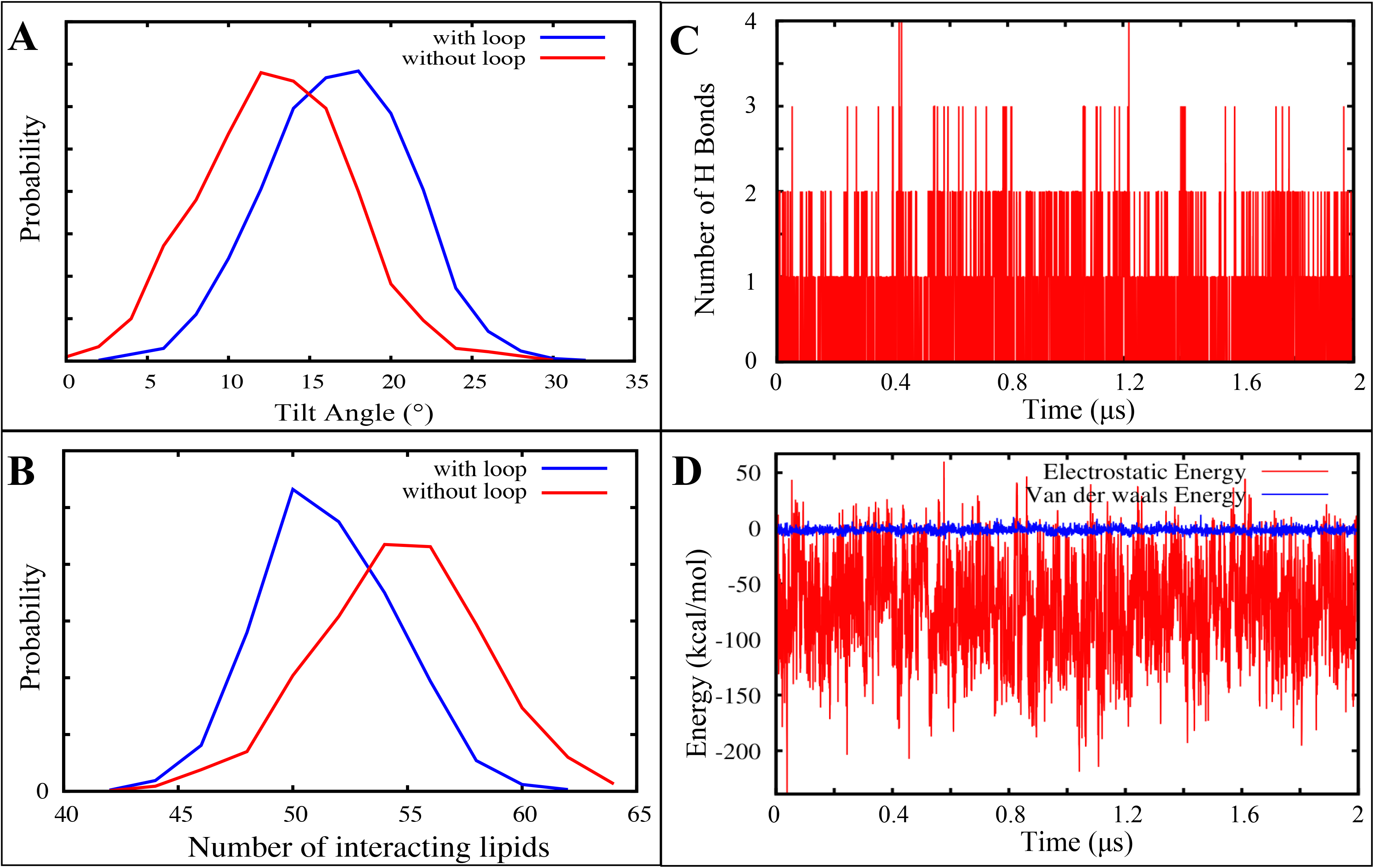
Influence of the C2 loop on membrane interactions. **A** The distribution of the tilt angle of YidC2 TM region with (blue) and without (red) the C2 loop, measured as the angle between the third principal axis (roll axis) of the TM region of protein and the membrane normal. **B** The distribution of the number of lipids interacting with the protein in the presence (blue) and absence (red) of the C2 loop. **C** Interactions between D207 of the C2 loop and the POPE lipid head groups measured by counting the number of hydrogen bonds between the two as a function of simulation time. **D** Electrostatic (red) and van der Waals interaction energies (blue) of C2 loop and lipids.

Phospholipids are specific participants in determining membrane protein organization^37^. We have recently shown that a slight change in the polarity of the head groups of a membrane’s constituent lipids could substantially change the structural dynamics of a transmembrane protein^31^. Specifically, the PE head groups were shown to play a key role in the structural dynamics of a bacterial ATP-binding cassette transporter^31^. The importance of the membrane composition has been shown extensively for various membrane proteins and other membrane-related phenomena^38,39,40,41^. It is thus reasonable to assume that the absence of the phospholipids in the crystallographic conditions could result in deviations of the resolved protein structure from its native conformation. This is also consistent with previous MD simulations of *E. coli* YidC, where the structure of the protein in a POPE containing lipid bilayer showed a more compact conformation compared with the crystallographic structure^25^.

Given the role of the lipid-protein interactions in the stability of the YidC2 protein with the C2 loop, it is important to test the reproducibility of our results in a lipidic environment that more closely resembles that of YidC2. We have thus simulated this protein, both with and without the C2 loop, in a heterogeneous membrane composed of 45% POPE, 45% 1-palmitoyl-2-oleoyl-sn-glycero-3-phosphoglycerol (POPG), and 10% cardiolipin (CL) for 240 ns (see Methods). This lipid composition resembles that of a native gram-positive membrane environment^42^. RMSD analysis clearly indicates that YidC2 is less stable when the C2 loop is absent in both the mixed lipid environment and the homogeneous POPE membrane. Figure 9A shows the RMSD of YidC2 with and without the loop in both membrane environments. Only the last 240 ns of the POPE simulations are used in this analysis for better comparison since the mixed lipid trajectories are only 240 ns. In each case, the reference structure for RMSD calculations is the average conformation over the entire 240 ns trajectory. The systems without the C2 loop show significantly more fluctuations than the systems with the C2 loop, irrespective of membrane type. Therefore, it is clear that the presence of anionic (POPG/CL) head groups in the lipid bilayer does not change our results regarding the role of C2 loop in global stabilization of YidC2. As shown in Figure 9B, the equilibrated C2 loop conformation is not significantly dependent on the presence or absence of the anionic lipids, at least within the timescale of our simulations.

**Figure 9.**
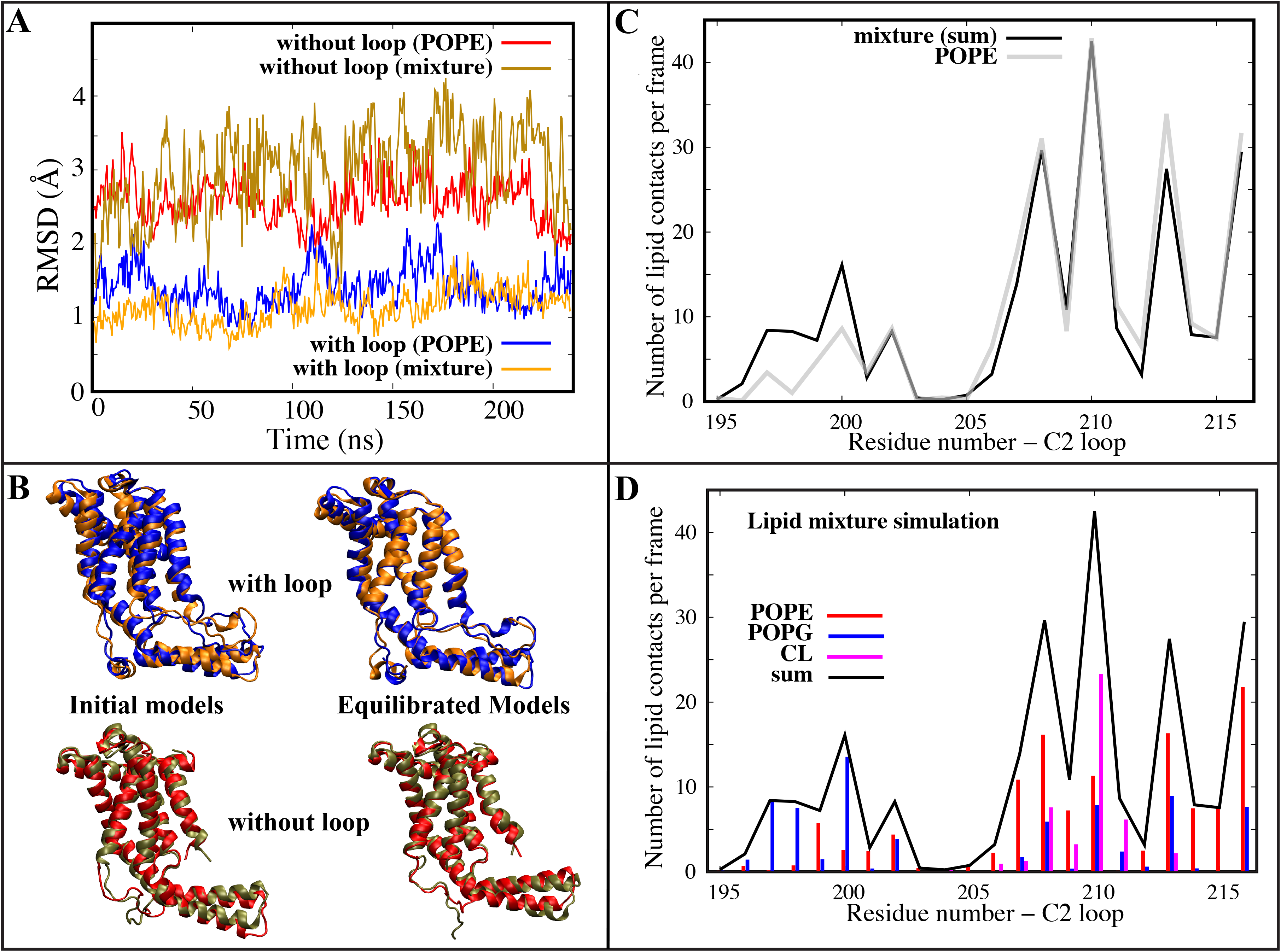
Behavior of YidC2 in a heterogeneous membrane environment. **A** RMSD time series of YidC2 from the last 240 ns of the 2-μs simulations in POPE with (blue) and without (red) the C2 loop and from the 240-ns simulations in mixed lipids with (orange) and without (bronze) the C2 loop. For each trajectory, the reference structure for RMSD calculations is an average structure from the 240-ns trajectory. **B** Cartoon representation of initial models and equilibrated conformations of YidC2 with the C2 loop, in POPE (blue) and mixed (orange) lipids, and without the C2 loop, in POPE (red) and mixed (bronze) lipids. **C** The number of contacts per frame between the C2 loop and lipids in homogeneous (grey) and heterogeneous (black) membranes. **D** The number of contacts per frame between the C2 loop and POPE (red), POPG (blue), and CL (magenta) lipids and their sum (black) in heterogeneous membrane simulations.

The salt-bridge interaction patterns observed in the POPE simulations (Figures 4 and S3) were also observed in the lipid mixture simulations (Supplemental Figure S7). In the system where the C2 loop is present, these include the inter-domain salt-bridge between D205 (C2 loop) and K109 (C1 region) and the slightly weaker interaction between D205 and K104 (C1 region) (Supplemental Figure S7A,B). A strong intra-domain interaction between E97 and R93 also occurs in the C1 region (Supplemental Figure S7C,D). The presence of these interactions in the lipid mixture simulations lends support to our proposal that salt-bridge interactions stabilize the C1 region when the C2 loop is present. In the system without the C2 loop, the intra-domain E97-R93 interaction is much weaker (Supplemental Figure S7E,F), which is similar to the interaction observed in the corresponding POPE simulation (Supplemental Figure S3C,D).

The carboxyl-terminus conformations are also similar in both systems with the C2 loop in the POPE and heterogenous membrane simulations (Figure 9B). The C-terminal helix remains helical only when the C2 loop is present (Supplementary Figure S8). Quantification and comparison of contacts between the C2 loop and the membrane reveals that the overall number of lipid contacts per frame is quite similar in both the POPE and lipid mixture simulations (Figure 9C). The only difference occurs in the region spanning residues 195-200, where the C2 loop preferentially interacts with POPG over POPE in the lipid mixture simulations (Figure 9D). However, when comparing the lipid-protein interactions of YidC2 in the presence and absence of the C2 loop, the same patterns are generally observed in both the POPE and heterogenous membrane simulations (Supplementary Figures S9 and S10).

The differential behavior of YidC2 in the presence and absence of the C2 loop is clearly reproducible in both a pure POPE membrane and a heterogenous POPE/POPG/CL membrane. We do not observe any significant difference between the system in the presence and absence of the anionic lipids POPG and CL. However, the only focus of this study is on the role of the C2 loop in protein dynamics in the inactive state of YidC2, assuming its crystal structure^5^ represents the inactive state. It is very likely that the anionic lipids such as POPG and CL could play an important role in other aspects of YidC2 dynamics and function.

Earlier we mentioned that the C2 loop was not resolved in the crystal structure of YidC2, where the crystals were grown in a lipidic cubic phase, using monoolein lipids^5^, which are quite different from phospholipids such as POPE, POPG, and CL. The absence of the phospholipids during the crystallization process of YidC2 could explain the fact that the C2 loop was not resolved. We propose that the absence of a physiologically relevant membrane in the crystallization process could cause the C2 loop appear disordered. We note that in all sets of simulations used in our study, we have used phospholipids with either anionic or zwitterionic head groups that are physiologically relevant and are very likely to contribute to the stability of the C2 loop in our simulations.

While our simulations clearly reveal the importance of the C2 loop in the stabilization of the YidC2 protein, it is unfortunate that this loop is not resolved in the crystal structure of YidC2 as discussed above. We note that the recently resolved *E. coli* YidC crystal structure contains a C2 loop, which is much shorter than that of the YidC2 C2 loop in gram-positive bacteria. The C1 and C2 loops in the resolved structure of *E. coli* YidC do not seem to be able to interact as in our YidC2 model. This observation is consistent with the fact that the two YidC proteins have significant functional differences.

To further assess the validity of our results, we also simulated YidC2 with an alternative C2 loop conformation in POPE for 240 ns, starting from the initial model of YidC2 used in the original 2-μs simulations of YidC2 in POPE. The initial conformations of the original and alternative C2 loops are 5.7 Å apart in terms of their RMSD (Supplemental Figure S10A) but the rest of the protein is identical. RMSD analysis indicates that YidC2 with the alternative C2 loop is just as stable as the protein with the original C2 loop (Supplemental Figure S11B). In the absence of the C2 loop, YidC2 is clearly destabilized in comparison to the systems with the original and alternative C2 loop conformations (Supplemental Figure S11B). The α-helical character of the C-terminus is retained throughout the alternative loop simulation (Supplemental Figure S10C). This is consistent with our results from the original C2 loop-POPE simulation (Figure 5B). Salt-bridge interactions observed in the original POPE simulations and lipid mixture simulations are also present in the POPE simulations with the alternative C2 loop conformation. The inter-domain salt-bridge between an aspartate residue in the C2 loop and a lysine residue (K109) in the C1 region is retained (Supplemental figure S12A,B). However, D207 is involved instead of D205. The strong intra-domain interaction between E97 and R93 in the C1 region is also retained (Supplemental Figure S12C,D).

We conclude that the second cytoplasmic loop of YidC2 could have a functional role perhaps by stabilizing the protein structure not only through its direct interactions with the C1 region and transmembrane helices TM4 and TM5 but also through its indirect effect on other transmembrane helices (particularly TM1) and the C-terminal region. The C2 loop of YidC2 is also significant due to its proximity to the periphery of the membrane on the cytoplasmic side and its strong interactions with the lipid head groups, a feature which is potentially absent in gram-negative YidC proteins. Due to its interactions with the membrane, the C2 loop was also found to change the tilt angle of the protein within the membrane. The C2 loop forms a salt bridge network with the functionally important C1 region and reduces its flexibility. The presence of the C2 loop also appears to reduce the flexibility of the carboxyl terminal region of the protein by increasing its helical propensity. Further research is needed to elucidate the importance of the C2 loop in the sec-independent insertion mechanism of small single-spanning membrane proteins such as the pf3 coat protein, whose interactions with the C1 region have been proposed to be crucial^5^. In the context of molecular dynamics simulations, our study suggests that modeling crystallographically unresolved loops may be necessary for accurate description of membrane protein dynamics, particularly if the missing loops are likely to interact with the periphery of the membrane.

## METHODS

We have used brute-force all-atom MD simulations to characterize the conformational transitions of bacterial YidC2 in a modeled membrane environment. We built two YidC2 systems; one with the C2 loop and another without the C2 loop. The crystal structure of bacterial YidC2 from *Bacillus halodurans* (PDB entry: 3WO7)^5^ with the missing C2 loop was initially processed using the Molecular Operating Environment (MOE)^43^ software to remove the crystal waters. MOE was also used to determine the appropriate protonation states for the titratable residues at physiological pH (7.4) using protonate3D facility of MOE, which resulted in standard protonation states, in agreement with the Propka 3.1^44^ predictions. For the system with the C2 loop, a Monte Carlo algorithm was used to model the missing C2 loop using the program Modeller^45^ to iteratively minimize the energy of the system. 10,000 Monte Carlo iterations were used to generate the C2 loop used as the initial model for the simulations of YidC2 with the C2 loop in POPE. The alternative C2 loop conformation, used in the control simulation of YidC2 in POPE lipids, was generated with 3,000 Monte Carlo iterations. The control simulations of YidC2 in the heterogenous lipids used the last frame of the equilibrated YidC2, with and without the C2 loop, from the simulations in POPE lipids (after 2 μs of simulations as described below). The CHARMM-GUI^46,47,48^ was then used to build the simulation systems. The protein was placed in lipids, solvated in a box of TIP3P waters, and 0.15 M NaCl (in addition to the counterions used to neutralize the protein) using CHARMM-GUI^46,47,48^. The systems with homogeneous POPE lipids consisted of 90/88 lipids in the upper/lower leaflet for the set with the C2 loop, and 90/96 lipids in the upper/lower leaflet for the set without the C2 loop. The system with the alternative C2 loop consisted of 94/89 lipids in the upper/lower leaflet. The heterogeneous membrane systems consisted of 45/45/10 POPE/POPG/CL lipids in the upper leaflet and 36/36/8 POPE/POPG/CL lipids in the lower leaflet. For the systems with the homogenous POPE lipids, the box size was ~80×80×102 Å^3^ with ~62,000 to 67,000 atoms in total. For the systems with the heterogeneous lipids, the box size was ~84×84×112 Å^3^ with ~ 75,000 atoms in total.

All systems were simulated with NAMD 2.10-13^49^ and the CHARMM36 all-atom additive force field^37,50^. Initially each system was energy-minimized for 10,000 steps using the conjugate gradient algorithm^51^. Then, we relaxed the systems by applying restraints in a stepwise manner (for a total of ~1 ns) using the standard CHARMM-GUI equilibration protocol^46^.

Production runs were performed for 2 μs for the systems with and without the original C2 loop in a POPE membrane. Production runs of simulation time 240 ns were performed for the system with the alternative C2 loop in a POPE membrane and the systems with and without the original C2 loop in a heterogeneous membrane (4.72 μs of simulation data in aggregate). The initial relaxation was performed in an NVT ensemble while all production runs were performed in an NPT ensemble. Simulations were carried out using a 2-fs time step at 310 K using a Langevin integrator with a damping coefficient of γ = 0.5 ps^−1^. The pressure was maintained at 1 atm using the Nosé-Hoover Langevin piston method^51,52^. The smoothed cutoff distance for non-bonded interactions was set to 10-12 Å and long-range electrostatic interactions were computed with the particle mesh Ewald (PME) method^53^. The trajectories were collected every 0.5 ns, resulting in 4,000 data points for each system for statistical analysis.

The TM helices and other subdomains were defined as follows: TM1 (63-83), TM2 (134-155), TM3 (175-190), TM4 (219-233), TM5 (233-258), C1 region (84 to 133), C2 loop (195 to 216), and modified C-terminal region (256 to 272). The last 13 C-terminal residues (268-280) were deleted from the wild-type sequence and replaced by a tag^5^. Residues 268 to 272 in the YidC2 model (modified C-terminal domain) used in our study belong to the tag.

The RMSD trajectory tool of VMD^54^ was used to calculate the RMSD and C_α_ atoms were considered for these calculations. For calculating the average RMSD, the entire trajectory was considered and error bars represent the standard deviation in the data. RMSF of individual residues was calculated using the *C_α_* atoms by aligning the trajectory against the crystal structure. The VMD timeline plugin^54^ was used to identify salt bridges and the cutoff distance used was 3.5 Å. The salt bridge plugin of VMD^54^ was used to calculate the distance between the two salt bridge residues over the course of the simulation, which is the distance between the oxygen atom of the participating acidic residue and the nitrogen atom of the basic residue. Lipid-protein interactions were characterized by counting the number of lipid molecules within 4 Å of the protein or any specific domain of the protein at every frame. PRODY software^55^ was used to carry out the PCA analysis. Only C_α_ atoms were used for PCA calculations. The VMD HBond plugin^54^ was used for hydrogen bond analysis; the cut-off distance and angles used were 3.5Å and 30° respectively. Dynamic network analysis was carried out using the dynamic network analysis tool in VMD^36^ and the program Carma^56^. In brief, the correlation of a residue pair *i* and *j* is defined as:

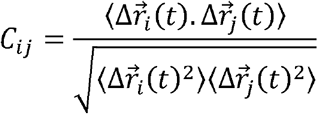

where 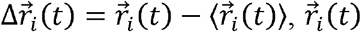 is the position of C_α_ atom of residue *i* at time *t*, and 〈·〉 is an average over all *t*. *C_ij_* quantifies the linear correlation of the motion of C_α_ atoms of residues *i* and *j*, with ±1 and 0 indicating the strongest positive/negative correlation and the complete lack of linear correlation, respectively. If and are measured under two different simulation conditions (e.g., YidC2 with and without the C2 loop), |*C_ij_* – *C′_ij_*| (the absolute value of the difference in the correlations between the two conditions) quantifies the absolute change due to the change in the condition (e.g., introduction/removal of the C2 loop in YidC2).

## ACKNOWLEDGMENTS

We thank the University of Arkansas, Fayetteville and the Arkansas Biosciences Institute for funding. This research was also supported by the Arkansas High Performance Computing Center, which was funded through multiple NSF grants and the Arkansas Economic Development Commission. This work also used the Extreme Science and Engineering Discovery Environment (XSEDE), which is supported by National Science Foundation grant number ACI-1548562. This work used Bridges at the Pittsburgh Supercomputing Center through XSEDE allocation MCB150129. We also thank Colin Heyes, James Walker, Mack Ivey, Adithya Polasa, and Mitchell Benton for helpful discussions.

## AUTHOR CONTRIBUTIONS

T.H., J.H., and M.M conceived the project and designed the simulations. T.H. conducted the simulations. T.H., J.H., S.H.T., V.G.K. and K.I. analyzed the data. T.H., V.G.K., K.I., and M.M. prepared the manuscript.

## AVAILABILITY OF DATA

The molecular dynamics trajectories and the analyses generated will be shared upon request to corresponding author.

