## Supporting Information for "The Role of a Crystallographically Unresolved Cytoplasmic Loop in Stabilizing the Bacterial Membrane Insertase YidC2"

Thomas Harkey[1], Vivek Govind Kumar[1], Jeevapani Hettige[1], Hamid Tabari[1], Kalyan Immadisetty[1], and Mahmoud Moradi[1*]

[1]Department of Chemistry and Biochemistry, University of Arkansas, Fayetteville, Arkansas 72701, United States

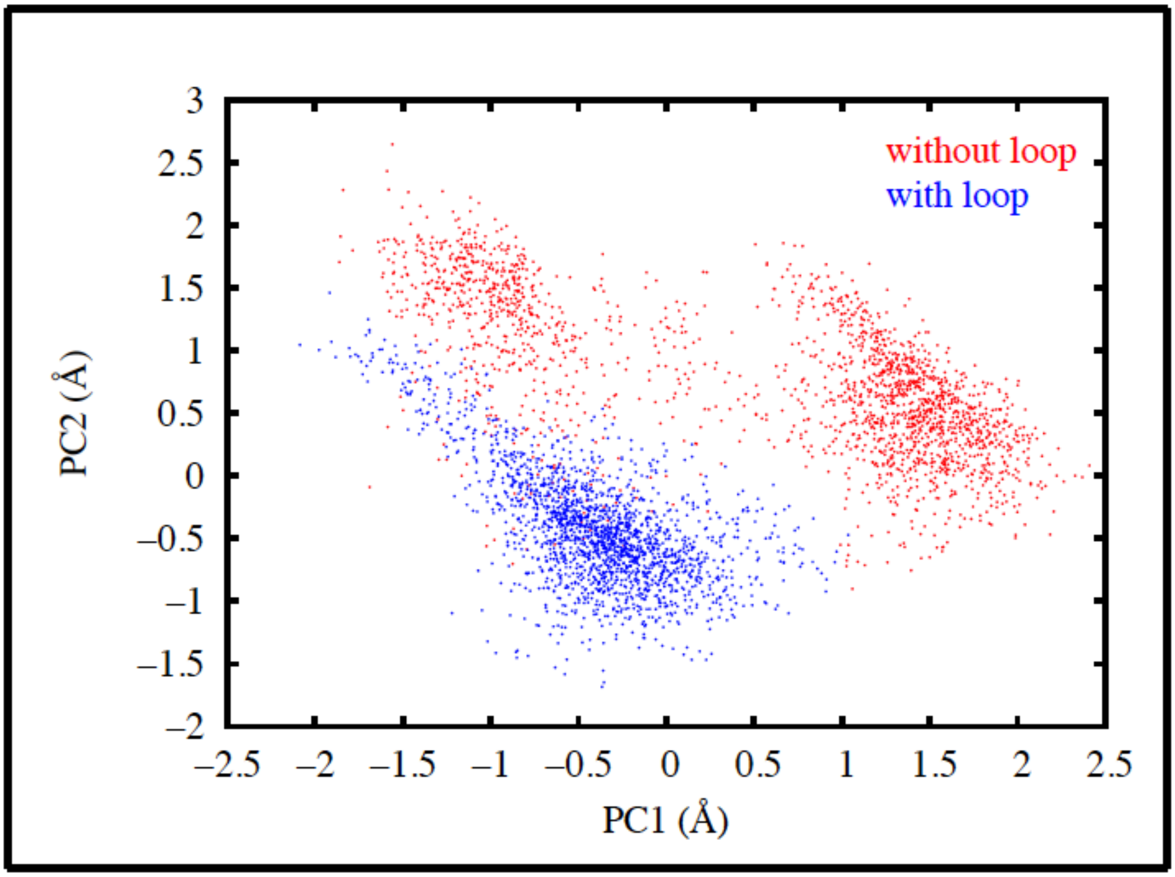


**Figure S1 Principal component analysis of the first microsecond of the YidC2 trajectories with and without the C2 loop (related to Figure 3).** Projection of the trajectories of YidC2 in POPE with (blue) and without (red) the C2 loop onto their first two principal components (PC1, PC2). The behavior of each system within the first μs is very similar to the overall result obtained from the 2μs trajectories (Figure 3). The YidC2 model with the C2 loop is quite stable and only fluctuates locally around its average structure. However, the model without the loop jumps between multiple conformations indicating a significant conformational flexibility. The dot product of the PC1 vectors from the two PCA analyses (one based on first half and one based on the entire trajectories) is 0.76, indicating a significant correlation. Similarly the dot product of the PC2 vectors is 0.75.

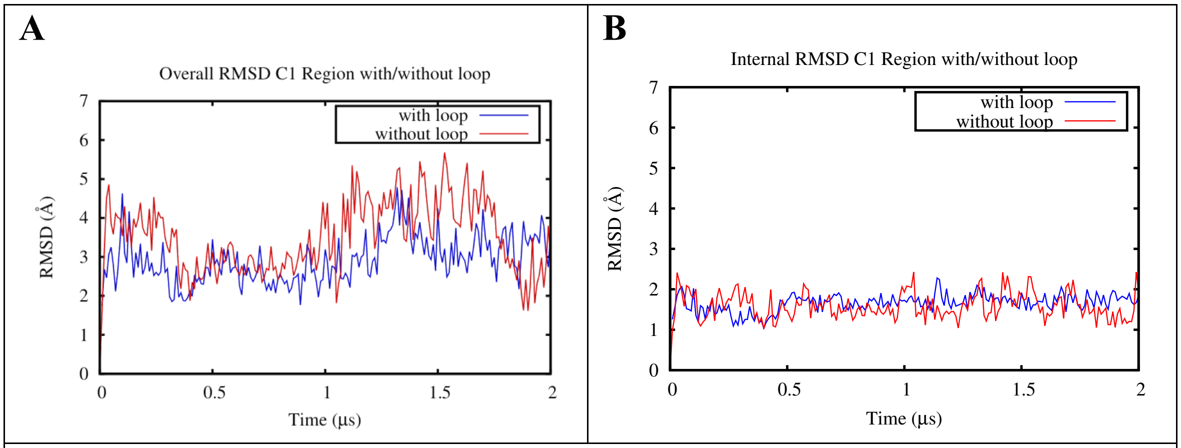


**Figure S2 C1 region stability assessed through the RMSD calculations (related to Figure 2). A** Overall and **B** internal RMSDs of the C1 region of the system with (blue) and without (red) the C2 loop, respectively. The internal RMSD does not show a meaningful difference between the two models but the overall RMSD shows a difference. This indicates the difference is mostly due to a rigid-body movement of the C1 region. For overall RMSD, the protein was aligned against the crystal structure and RMSD was calculated for the region of interest (here, C1 region) with respect to its initial configuration. For internal RMSD, we aligned the region of interest (here, C1 region) against its own initial configuration and calculated RMSD with respect to this configuration.

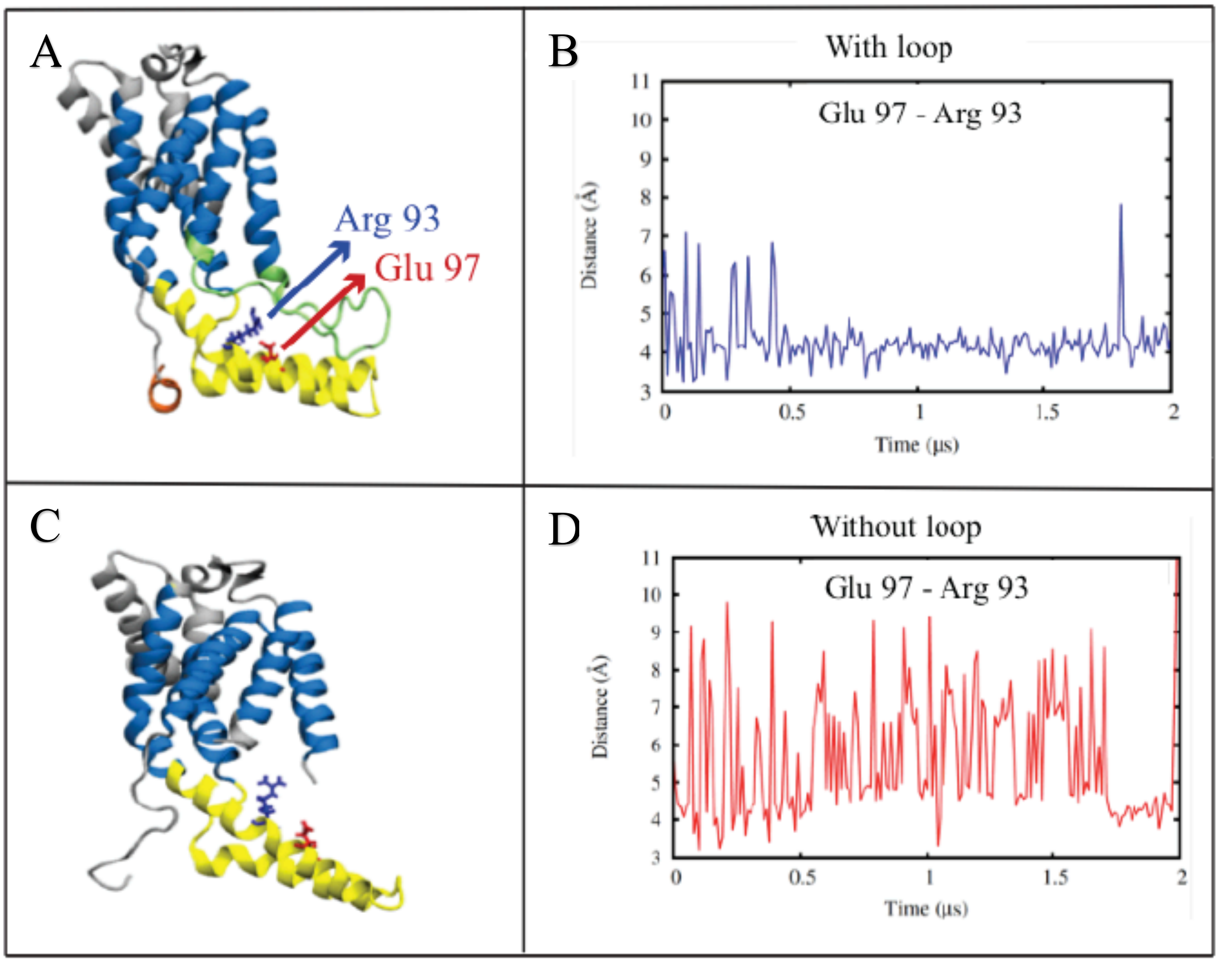


**Figure S3 C1 region intradomain salt bridge formation (related to Figure 4). A** An intradomain salt bridge is formed between Glu 97 and Arg 93 in the model with the C2 loop. **B** Time series associated with the average donor-acceptor distance of Glu 97 and Arg 93 residues in the model with the C2 loop. **C** The salt bridge between Glu 97 and Arg 93 is rarely formed in the model without the C2 loop. **D** Time series associated with the average donor-acceptor distance of Glu 97 and Arg 93 residues in the model without the C2 loop.

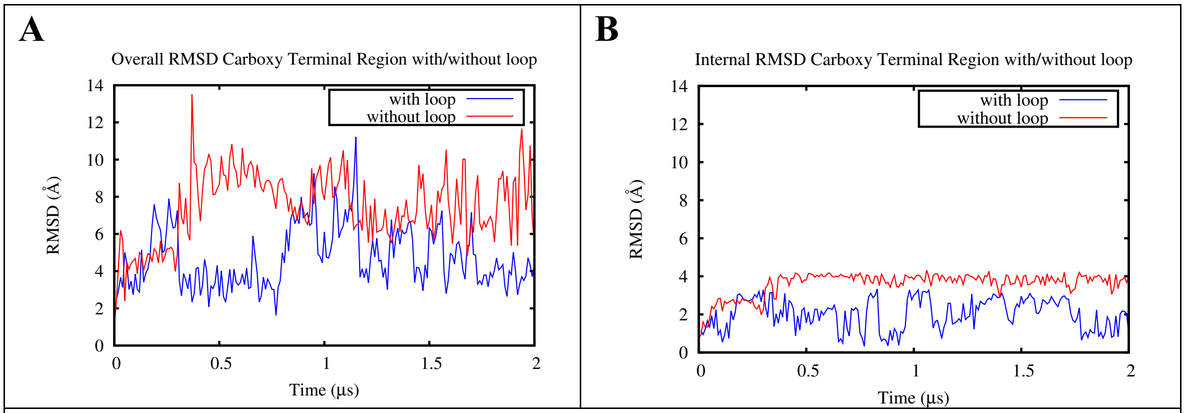


**Figure S4 RMSD time series of the carboxyl terminus (related to Figure 5). A** Overall and **B** internal RMSDs of the C-terminus of the system with (blue) and without (red) the C2 loop, respectively. See Figure S2 for the definition of overall and internal RMSDs. The difference between the two models is not limited to the rigid-body movement of the C-terminus but the internal conformation of the C-terminal domain is also different in the two models.

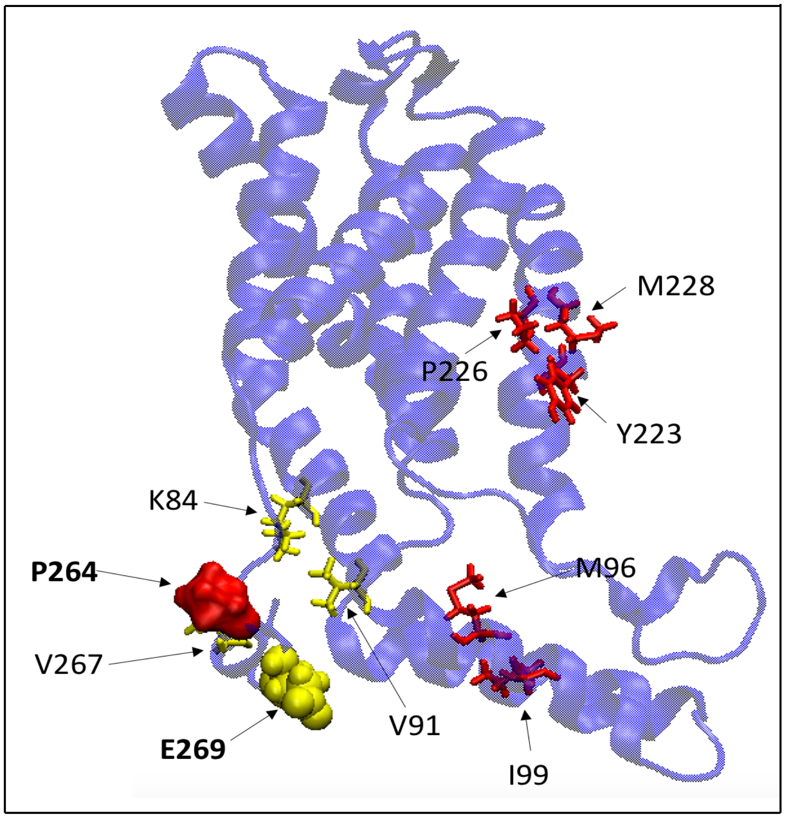


**Figure S5 Cartoon representation of the residues with the most significant cross-correlation changes due to the presence/absence of the C2 loop (related to Figure 6).** The largest changes are observed for C-terminal residues P264 (correlating with C1 region residues such as K84 and V91) and E269 (correlating with C1 region residues such as M96 and I99 and TM4 residues such as Y223, P226, and M228), colored red and yellow respectively.

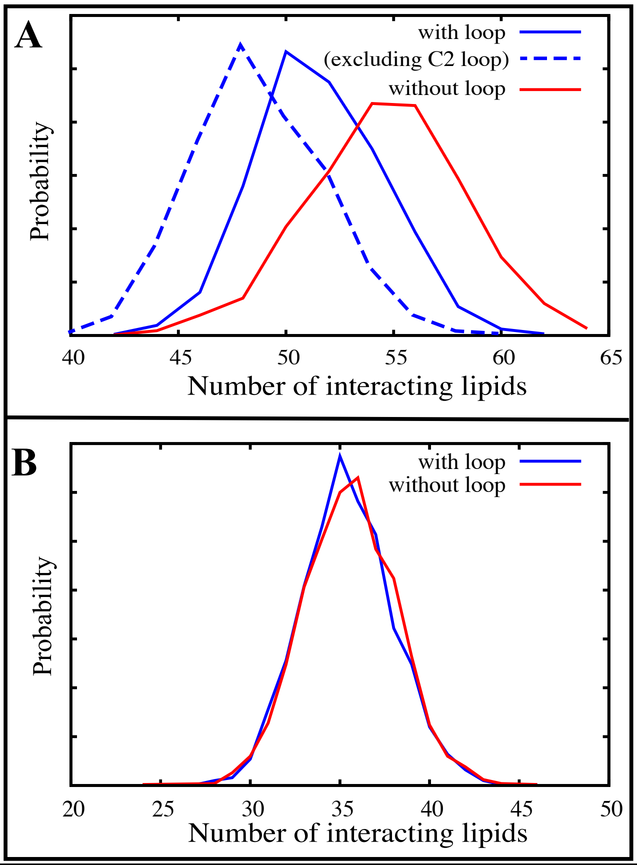


**Figure S6 Lipid-protein interactions in YidC2 simulations with and without the C2 loop (related to Figure 8). A** Same as Figure 8B, also showing the number of lipids interacting with protein per frame excluding the C2 loop from calculations for the YidC2 simulation with the loop (dashed line). The difference between the dashed and solid blue lines is due to the direct C2-lipid interactions. **B** Same as Figure 8B but only including the lipids that are directly interacting with the TM region of YidC2.

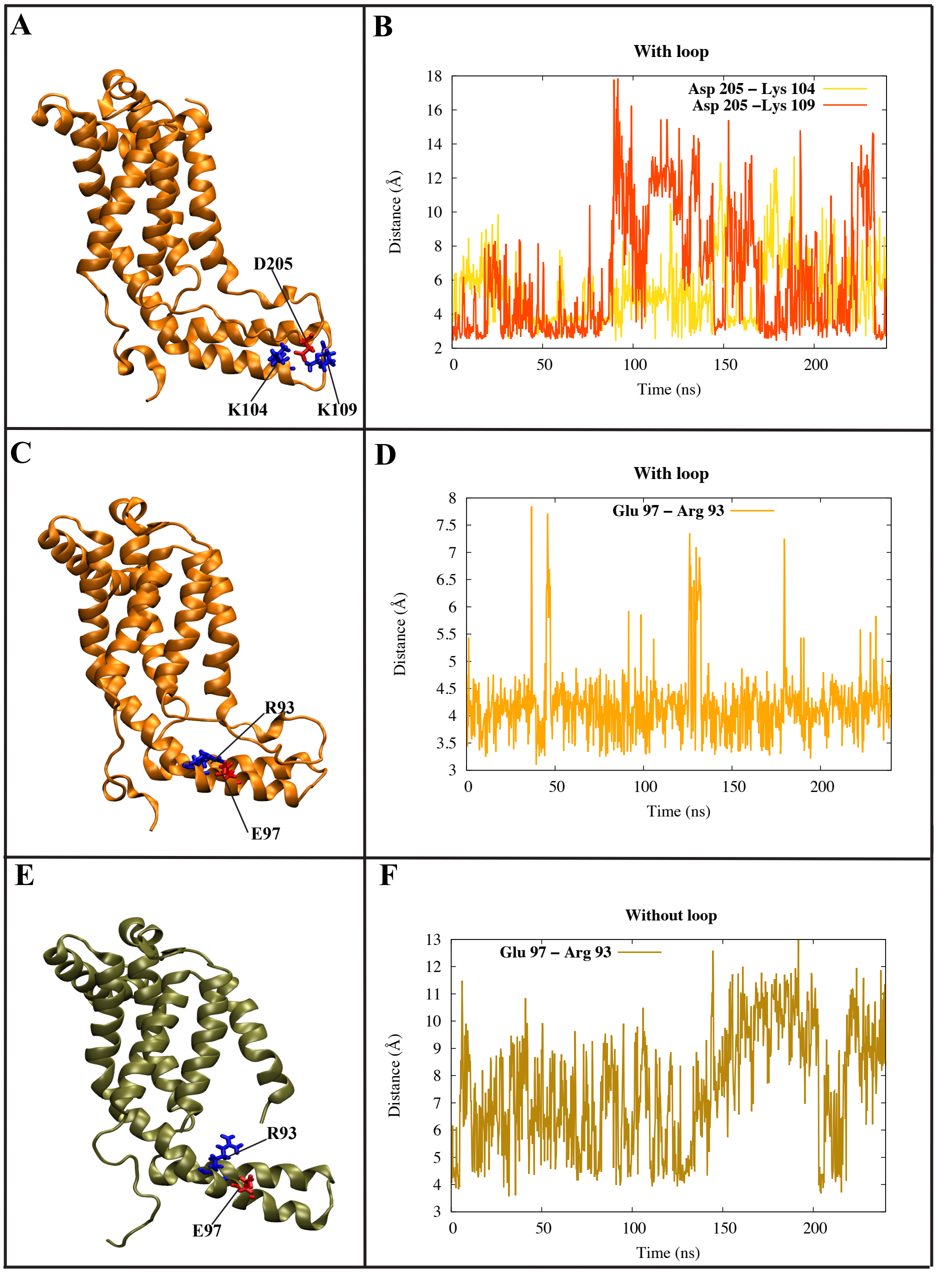


**Figure S7 Inter- and intra-domain salt bridge formation in the lipid mixture simulations (related to Figures 4 and 9). A** D205 of the C2 loop can potentially form an inter-domain salt bridge with K109 and/or K104 of the C1 region in the system with the C2 loop. **B** Time series of the D205-K109/104 donor-acceptor salt bridge distances. **C** An intra-domain salt-bridge is formed between E97 and R93 of the C1 region in the model with the C2 loop. **D** Time series of the E97-R93 donor-acceptor distance in the model with the C2 loop. **E** The salt bridge between E97 and R93 is rarely formed in the model without the C2 loop. **F** Time series of the E97-R93 donor-acceptor distance in the model without the C2 loop.

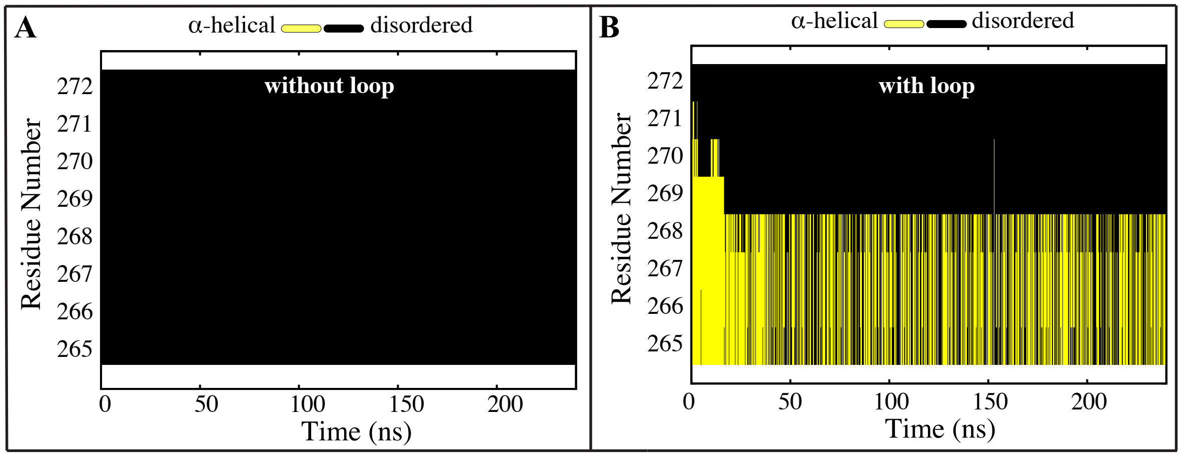


**Figure S8 α-helical character of the c-terminus in the lipid mixture simulations (related to Figure 5).** *α*-helical character of the residues 265 to 272 of the modified carboxyl terminal domain **A** without and **B** with the original C2 loop in the lipid mixture simulations is shown as a function of the simulation time. When the loop is absent, the C-terminal region unravels into a coil. When the loop is present, the *α*-helical character of the C-terminus is retained throughout the 240 ns simulation.

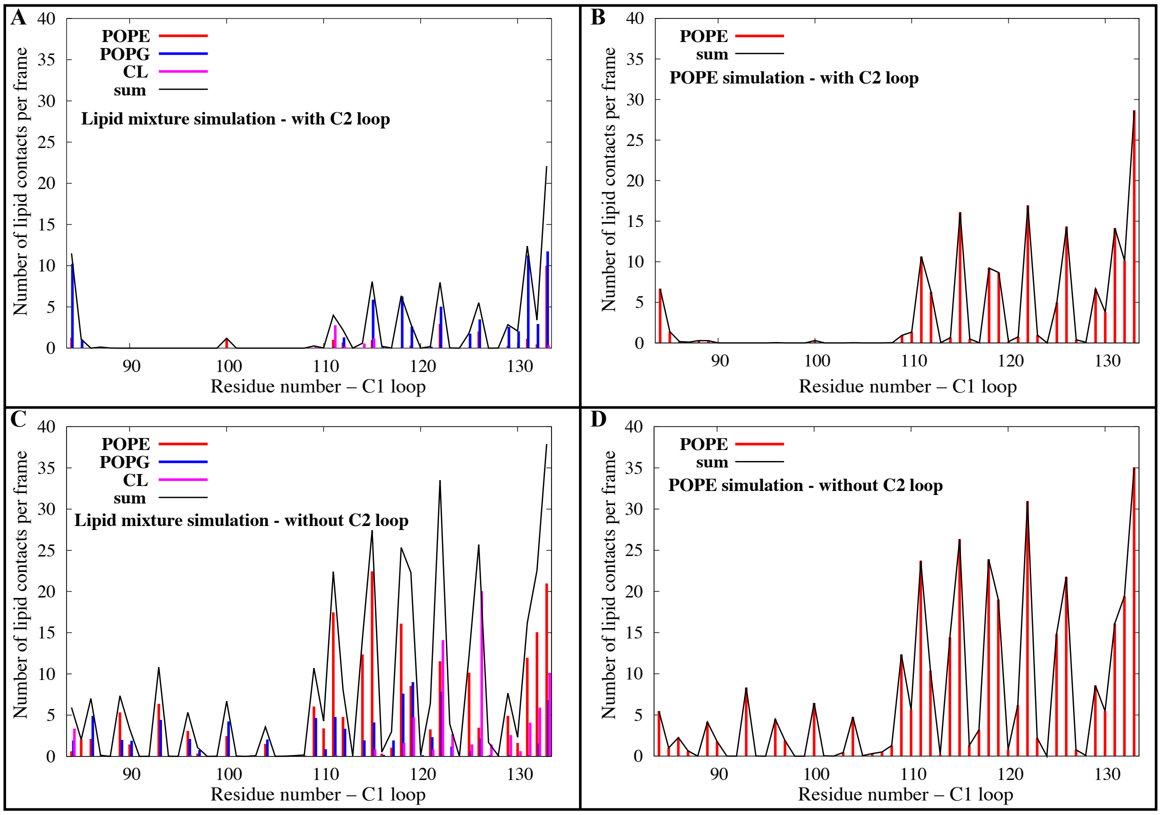


**Figure S9 Quantification of C1-lipid interactions in POPE and lipid mixture simulations of YidC2 with and without the C2 loop (related to Figure 9).** The number of contacts per frame between the C1 loop and POPE (red), POPG (blue), and CL (magenta) lipids and their sum (black) in simulations of **A** YidC2 with the C2 loop in a heterogeneous membrane, **B** YidC2 with the C2 loop in a homogeneous membrane, **C** YidC2 without the C2 loop in a heterogeneous membrane, and **D** YidC2 without the C2 loop in a homogeneous membrane.

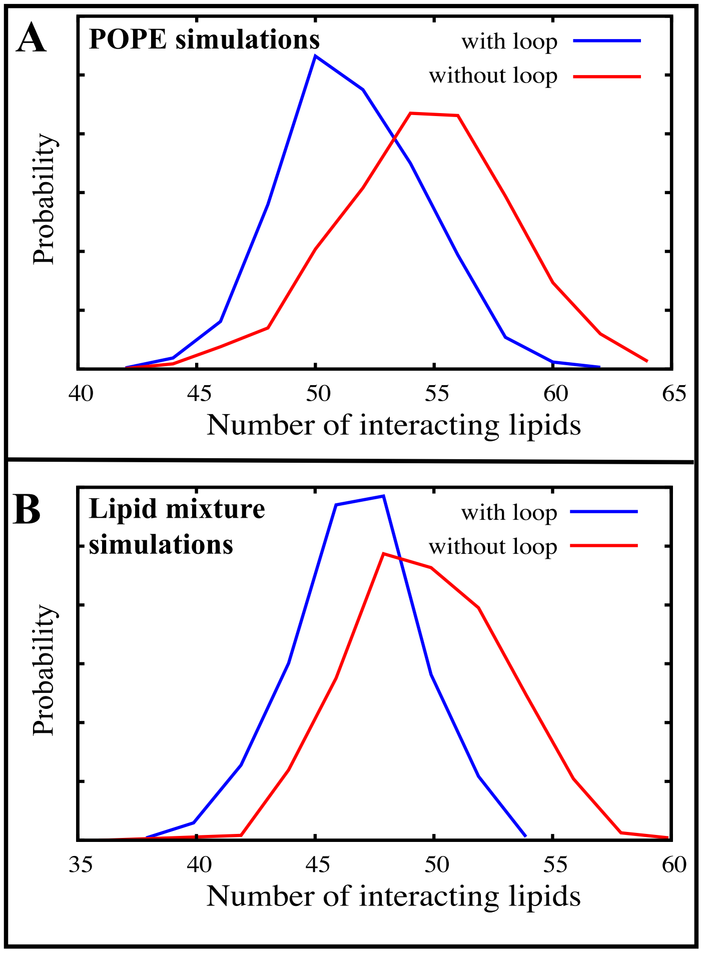


**Figure S10 Comparison of lipid-protein interactions in POPE and lipid mixture simulations of YidC2 with and without the C2 loop (related to Figures 8 and 9). A** Panel B of Figure 8, repeated here for the ease of comparison, which shows the distribution of the number of lipids interacting with YidC2 based on POPE simulations with and without the C2 loop. **B** Same as Figure 8B but based on lipid mixture simulations.

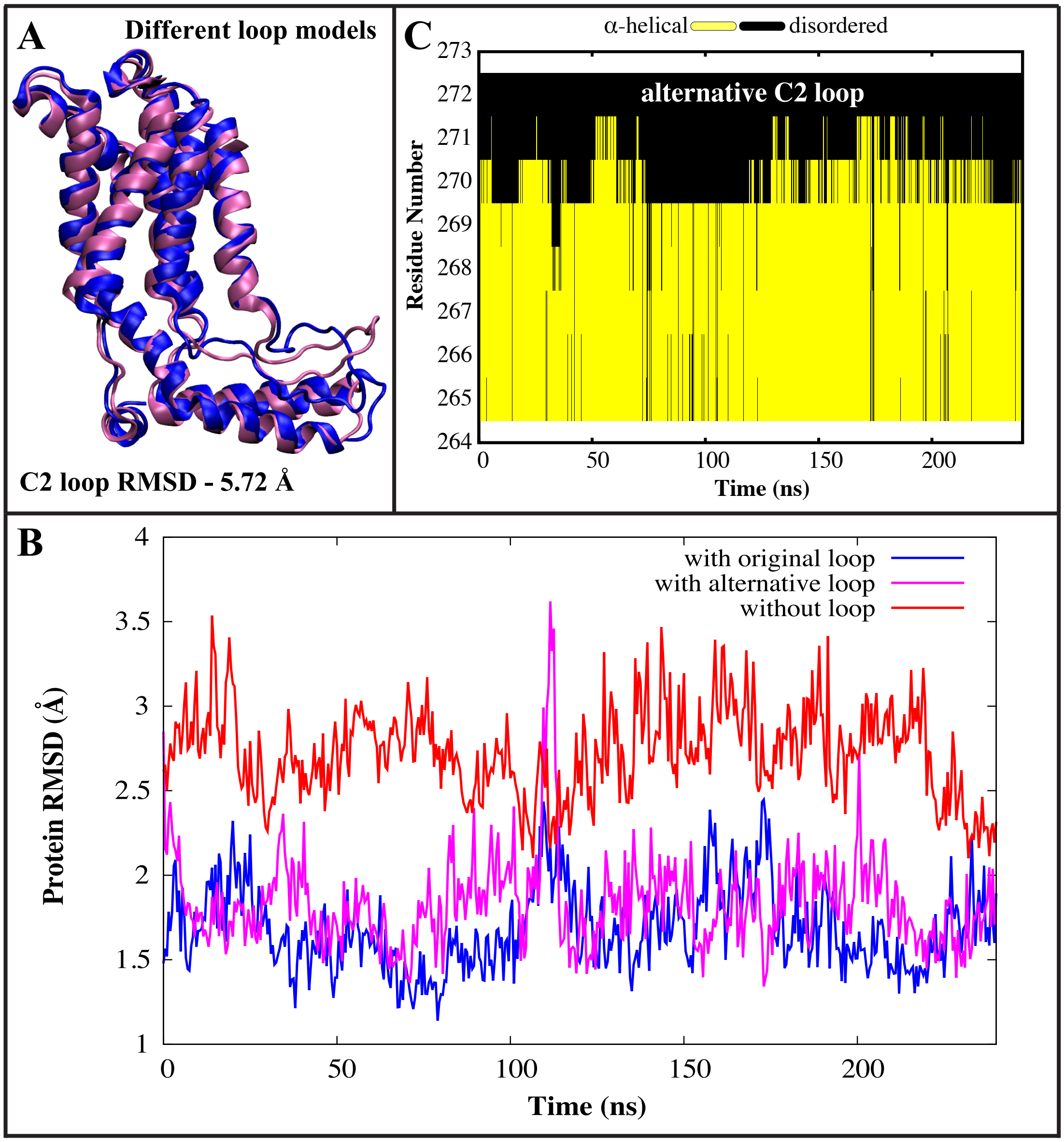


**Figure S11 Behavior of YidC2 with an alternative C2 loop conformation in POPE (related to Figures 2 and 5). A** Comparison of initial models of YidC2 with the original C2 loop (blue) and with the alternative C2 loop conformation (magenta). Initial models were obtained from the first frames of the original 2-μs POPE simulation and the 240 ns alternative C2 loop-POPE simulation respectively. **B** Comparison of the RMSD time series for YidC2, with the original (blue) and (magenta) alternative C2 loops, and without (red) the C2 loop in POPE membrane. For the original C2 loop simulations and those without the C2 loop, we have used the last 240 ns of the 2-μs simulations. The reference structure for RMSD calculations is the average conformation of the 240-ns trajectory shown in each case. Both conformations of the C2 loop stabilize YidC2. **C** α-helical propensity of the carboxyl terminal domain of YidC2 with the alternative C2 loop is shown as a function of the simulation time. The α-helical character of the C-terminus is retained throughout the 240 ns simulation.

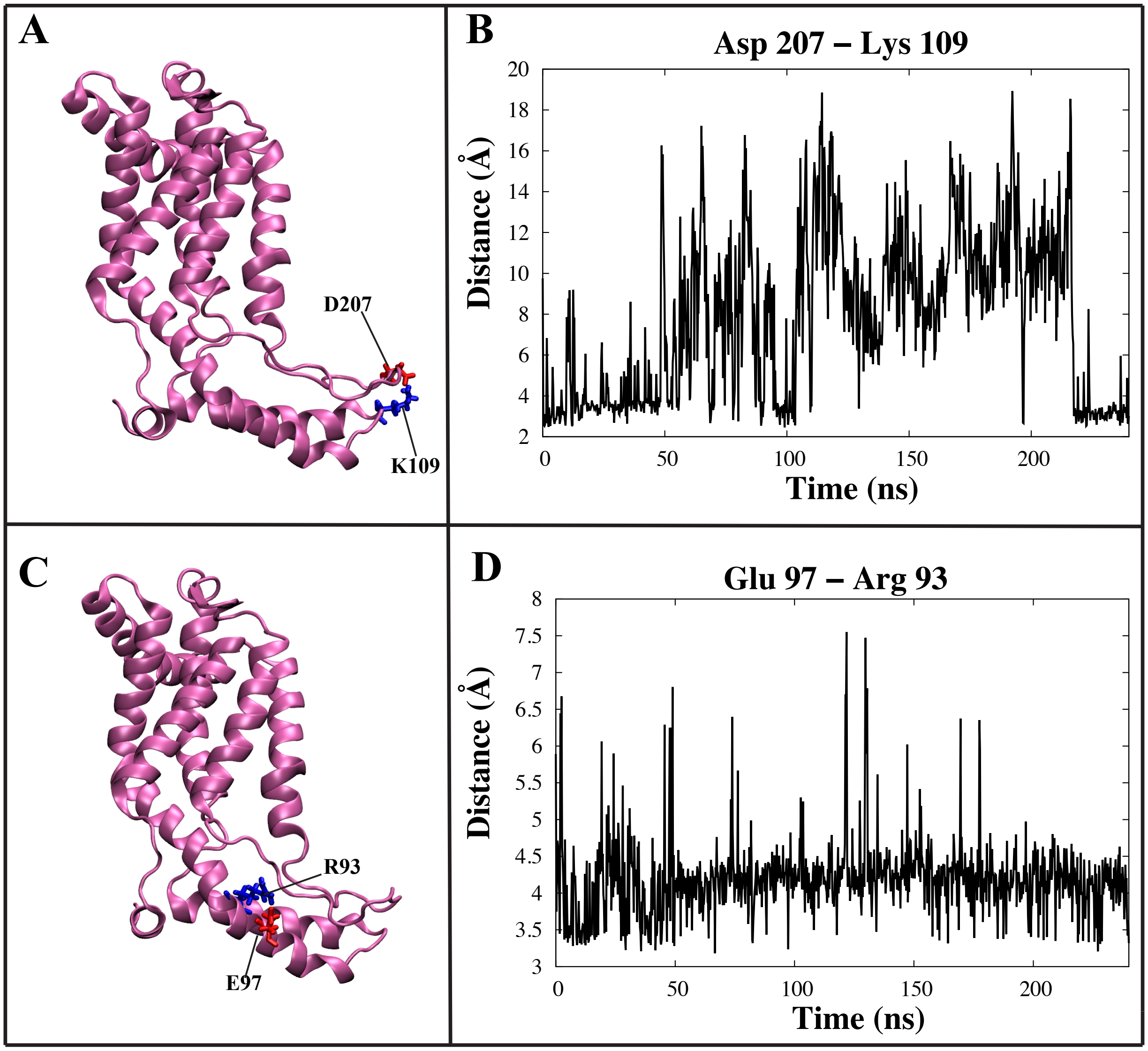


**Figure S12 Inter- and intra-domain salt bridge formation in the alternative C2 loop-POPE simulations (related to Figure 4). A** D207 of the C2 loop forms an inter-domain salt bridge with K109 of the C1 region. **B** Time series of the D207-K109 donor-acceptor salt bridge distance. **C** An intra-domain salt-bridge is formed between E97 and R93 in the C1 region. **D** Time series of the E97-R93 donor-acceptor distance.
